# Exceptionally high sequence-level variation in the transcriptome of *Plasmodium falciparum*

**DOI:** 10.1101/2021.05.14.444266

**Authors:** Bruhad Dave, Abhishek Kanyal, DV Mamatharani, Krishanpal Karmodiya

## Abstract

Single-nucleotide variations in RNA (hereafter referred to simply as SNVs), arising from co- and post-transcriptional phenomena including transcription errors and RNA editing, are well studied in organisms ranging from bacteria to humans. In the malaria parasite *Plasmodium falciparum*, stage-specific and non-specific gene-expression variations are known to accompany the parasite’s array of developmental and morphological phenotypes over the course of its complex life cycle. However, the extent, rate and effect of sequence-level variation in the parasite’s transcriptome are unknown. Here, we report the presence of pervasive, non-specific SNVs in the transcriptome of the *P. falciparum*. We show that these SNVs cover most of the parasite’s transcriptome. SNV rates for the *P. falciparum* lines we assayed, as well as for publicly available *P. vivax* and *P. falciparum* clinical isolate datasets were of the order of 10^−3^ per base, about tenfold higher than rates we calculated for bacterial datasets. These SNVs may reflect an intrinsic transcriptional error rate in the parasite, and RNA editing may be responsible for a subset of them. This seemingly characteristic property of the parasite may have implications for clinical outcomes and the basic biology and evolution of *P. falciparum* and parasite biology more broadly, and we anticipate that our study will prompt further investigations into the exact sources, consequences and possible adaptive roles of these SNVs.

## Introduction

Fidelity in the transcription of DNA into RNA and the correct translation of mRNAs into proteins is crucial. Accurately made proteins produce “correct” phenotypes and ensure that the cell survives. DNA mutations represent a significant source of variation in the flow of genetic information, and many affect phenotypes and become established as single-nucleotide polymorphisms (SNPs) in a given population of cells. For unicellular parasites, these SNPs are a conceivably essential way to adapt to life in their respective hosts over many generations. Proteins are largely immutable forms of genetic information, but the transcriptional landscape provides much scope for diversification.

A previous body of work has shown an inherent heterogeneity in the gene-expression levels as well as copy-number variation^**[5]**^ in *P. falciparum*, a parasite that still affects over 200 million people worldwide^**[10]**^. These observations have been reported across multiple conditions and levels -- in untreated parasite cultures^**[6]**^ and as a response to physiological-like stressors^**[7]**^; at the population^**[8]**^ and the single-cell^**[9]**^ level, and between clinical isolates and lab-adapted cultures^**[5]**.^ Antigenic variation in *P. falciparum* is well-documented^**[11]**^, and the parasite exhibits alternative splicing^**[12,13,14,15]**^; such transcriptional variation in essentially clonal populations represents another potential layer of complexity that is likely to affect clinical outcomes^**[16,17]**^, and studies have shown that heterogeneity through gene expression variation serves as a population-level survival strategy in unicellular organisms^**[18,19]**^

Sequence-level variation -- single-nucleotide variations (SNVs) and insertion-deletion events (indels) -- is another potential source of population-level transcriptomic diversity. In the form of transcriptional error rates and RNA editing, it has been extensively described in systems including bacteria^**[1,2]**^, yeast^**[20]**^, C. elegans^**[21]**^, cephalopods^**[22]**^, rodents^**[23,24]**^, and humans^**[3,4]**^. However, studies on such heterogeneity in the *P. falciparum* transcriptome are largely missing and such variations have the potential to impact downstream sources of transcriptional heterogeneity, which may facilitate stress adaptation and drug-resistance generation. To investigate the extent and rate of SNVs in *P. falciparum*, we analysed transcriptomic data from a lab-adapted parasite culture (strain 3D7), to obtain three datasets - an untreated control, a temperature-stressed culture and a drug-stressed culture. We also analysed RNAseq data from three drug-resistant *P. falciparum* lines sourced from the MR4 repository, namely MRA_1236, MRA_1240 and MRA_1241. We also performed whole-genome sequencing corresponding to each of the six resulting datasets, which allowed us to accurately discard transcriptomic sequence variations arising from genomic SNPs, without the need to use predictive or consensus-based methods. We then used REDItools 2.0^**[25]**^ to perform empirical variant-calling against the *P. falciparum* reference genome (v.41), and we found SNV rates on the order of 10^−3^ per base, a metric that was consistent across all six P. falciparum samples. In this work, we describe the spectrum of base substitutions and their predicted functional effects.

## Results

### A Global View of Transcriptional Sequence-Variation

We applied REDItools 2.0 (a tool originally designed to detect RNA-editing) to both the RNAseq data and the WGS data for each sample. We removed SNVs arising from genomic single nucleotide polymorphisms (SNPs) as well as discarded positions where RNAseq data showed no variation. Additionally, we ascertained optimum cutoffs for WGS coverage (**Supplementary Figure S1, Supplementary Table S3)**, RNAseq coverage (**Supplementary Figure S2, Supplementary Table S4)** and frequency of variation (the number of reads supporting a variant nucleotide divided by the total number of reads covering that position) (**Supplementary Figure S3, Supplementary Table S5)**. SNVs not supported by at least 10 WGS reads and 5 RNAseq reads were removed. Finally, we retained only those SNVs whose frequency of variation was greater than or equal to 0.1. In our estimation, this combination of cutoffs ensures that sequencing errors and other technical errors are largely filtered out, and mitigates any technical variability. We noted that the rates of occurence of SNVs -- which were spread out all over the genome (**Figure 1A**) -- were not dependent on sequencing and alignment statistics of the sample, indicating that they are consequences of biological characteristics of the parasite, rather than technical properties of the sequencing runs and alignment methods (**Supplementary Table S1**).

**Figure 1:**
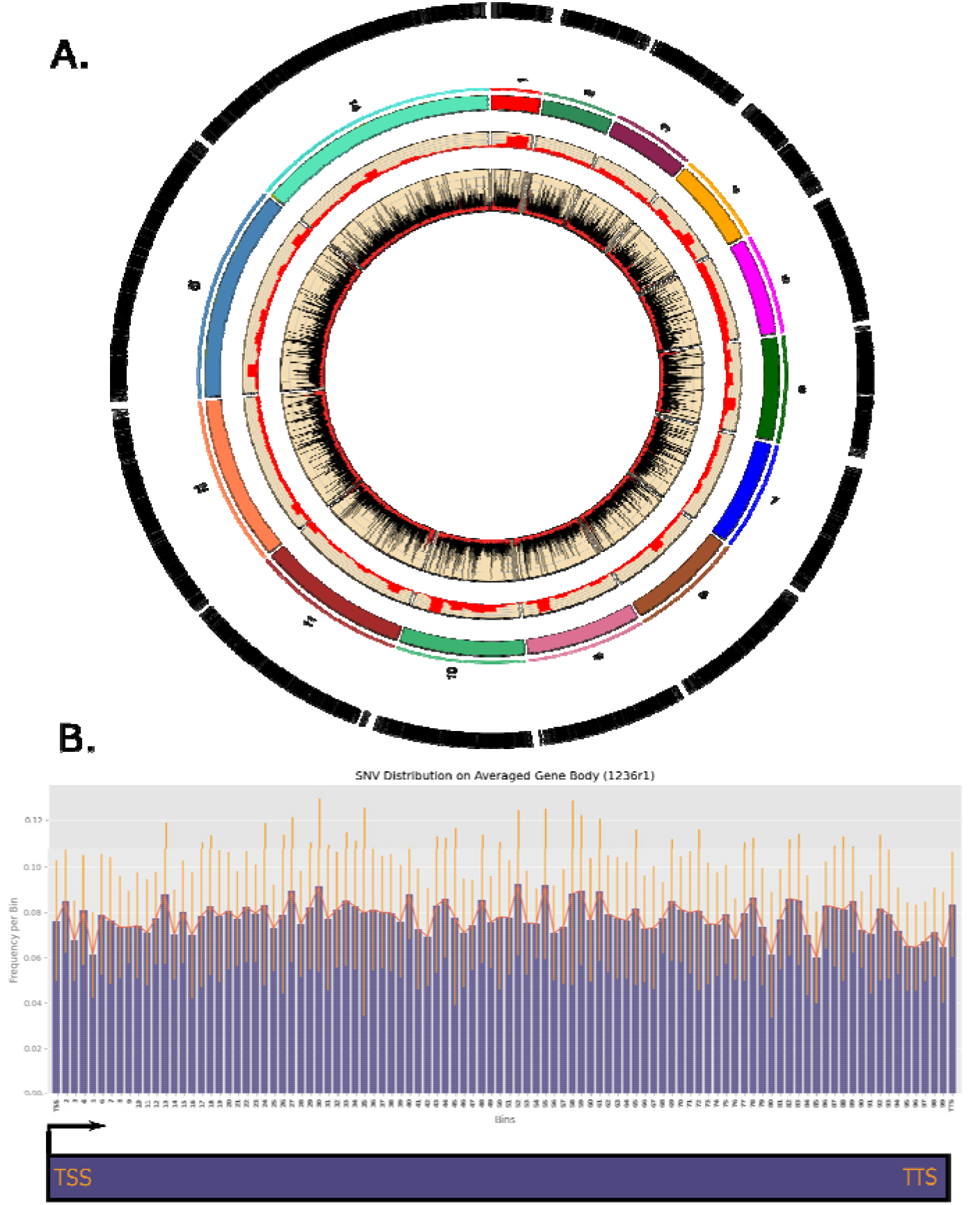
A Global View of Single-nucleotide Variations in Plasmodium falciparum. (data shown for a representative sample, MRA_1236 replicate 1). Plot showing the pervasive nature of SNVs. From outward: Track 1: SNV positions represented as vertical lines; Track 2: Ideogram of chromosomes, proportional to chromosome lengths; Track 3: Histogram of RNAseq coverage at recorded positions; Track 4: Area plot of frequency of variation at recorded positions. Histogram showing distribution of SNVs on an averaged gene body.

Following filtering, each sample showed ~3×10^4^ variant positions (**Supplementary Table S1**). Annotating SNVs with the genes in which they occurred showed that they were located all across the transcriptome, affecting ~3800 of 5700 genes (**Supplementary Table S2**) in all samples. To check whether the SNVs were biased toward either end of the transcript, we constructed an average histogram of SNV frequency, dividing each affected gene into bins of equal width, and then assigning bins to each SNV. We observed that SNVs occupied the whole averaged gene length and did not show a specific positional bias along the gene body (**Figure 1B**). Notably, replicate datasets for a given strain showed only a small amount of overlap, further indicating the pervasive, non-specific nature of the SNVs (**Supplementary Figure S4**). We also noted that the relative frequency of occurrence of SNVs for a given gene (defined as the number of SNVs found in that gene divided by the length of that gene) showed only a very small negative correlation or no correlation to the expression levels (in transcripts per million) of that gene, the outlier being one replicate dataset for MRA_1240. (**Supplementary Figure S5**).

### The Spectrum of SNV Types and Effects

In order to understand the variations in greater detail, we characterised the range of nucleotide substitutions and their probable functional effects. We observed that A-to-G changes and T-to-C changes predominated, each representing 23-27% of the total number of SNVs found (**Figure 2A**). A-T and vice versa substitutions (~14% each, of the total number of SNVs) were the second most abundant on average. G-to-C and vice versa substitutions were the least abundant SNV type (**Figure 2A, Supplementary Table S6**). These proportions were relatively well-conserved between the drug-resistant MRA lines as well as the drug-sensitive 3D7 culture. To check whether these base-change characteristics depended on environmental stresses, we subjected the 3D7 *P. falciparum* 3D7 line to temperature stress (at 40□) and separately, to mild drug stress (dihydroartemisinin at 30 nM), each for six hours at the early trophozoite stage. However, these stresses had a minimal effect on the base-change profile, and the proportions of nucleotide substitutions were remarkably similar between the control *P. falciparum* 3D7 culture and the two stressed cultures (**Figure 2A**). With the assumption that all base substitutions are equally probable, we performed a chi-squared test and found that the frequency distribution of base substitutions we observed differed significantly from the expected uniform distribution (p = 3.0147×10^−200^ □ ^2^ = 1434.7, df = 167). Interestingly, while the base-change pattern showed that most of the nucleotides that changed changed (i.e. reference/focal nucleotides) were As and Ts, the probabilities of any reference nucleotide changing to a G or a C were only slightly greater than those of a focal nucleotide changing to an A or a T (**Figure 2B, Supplementary Table S7**). The former observation is an expected outcome of the AT-richness of the *Plasmodium* genome. However, the fact that any given reference nucleotide changed to A, T, G or C in very similar proportions suggests that, if the SNVs we observe arise from RNA Polymerase II (RNA Pol II) errors, then the enzyme has no specific tendency toward mis-incorporating a given nucleotide. Given the bias of the spectrum of base-substitutions toward the A-to-G and T-to-C substitution types, we speculated that RNA editing may be responsible for a subset of SNVs. We tested this hypothesis by searching for sequence-motifs centered on or around SNVs as well as performing a BLAST-based search for potential RNA-editing enzymes in the proteome of the parasite. We used human and *Trypanosoma brucei* enzymes known to be RNA-editors as query sequences for the latter analysis, but we did not find any high-confidence candidate RNA-editors in *P. falciparum*, nor any conserved motifs (**Supplementary Table S8)**.

**Figure 2:**
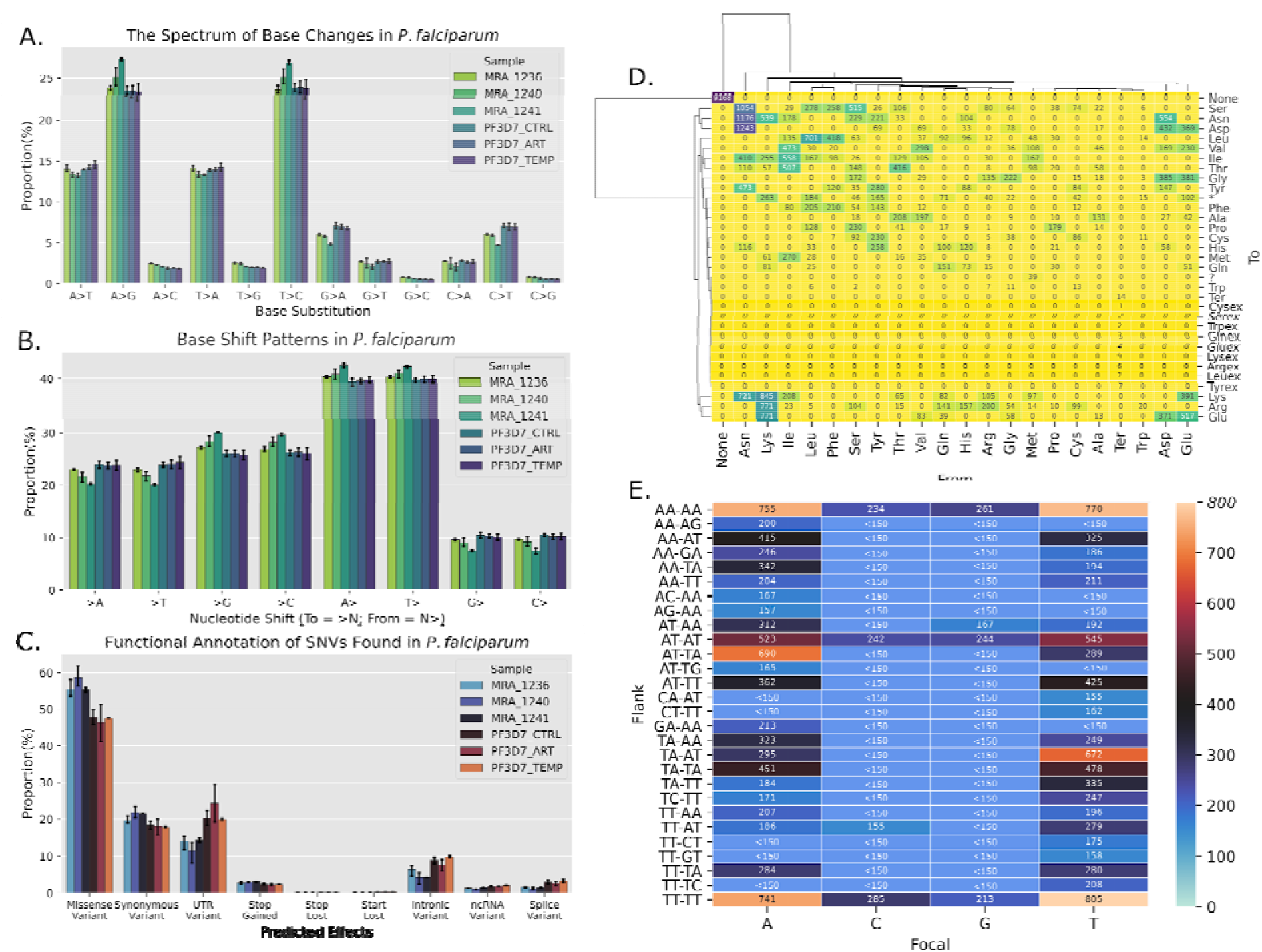
SNV Types and Effects. (A) Base changes (Percentage of Total) (B) Base shift patterns (%X> denotes the proportion of SNVs where base X changed to another base; %>X denotes the proportion of SNVs where the reference base was X). The distribution of predicted functional effects: (D) Representative heatmap of the spectrum of amino acid changes. (E) Representative heatmap of the abundance of the most common dinucleotide patterns flanking each focal (original/reference) base. (A), (B), (C): Error bars represent 95% confidence intervals; number of replicates = 2 (3D7_CTRL, 3D7_ART, 3D7_TEMP, MRA_1241), 3 (MRA_1236, MRA_1240) (D) and (E) show data from sample MRA_1236 replicate 1.

Further, we annotated effects to the SNVs using snpEff^**[26]**^ to investigate the range of predicted functional consequences, and filtered the snpEff outputs using snpSift^**[27]**^ to analyse amino acid change patterns. A small percentage were found in intronic regions, possibly reflecting alternative splicing in some transcripts (**Figure 2C, Supplementary Table S9**). The majority of SNVs in the coding region were missense (47-60%), with about 20% of them being synonymous, and about 4% being nonsense changes (**Figure 2C**). These proportions were also well conserved between all three MRA lines, the 3D7 control and two 3D7 stressed cultures. Asparagine was consistently the most changed amino acid, with lysine, isoleucine, aspartate, glutamate, leucine, phenylalanine, serine and tyrosine rounding out the most changed residues, although a proportion of said amino-acid shifts were like to like, that is, synonymous codon changes (**Figure 2D, Supplementary Figure S6**). We also tested whether focal nucleotides had characteristic flanking sequences that might increase their propensity to be changed. To this end, we extracted a pentanucleotide sequence for each variant position, taking two nucleotides on either side of each focal nucleotide, and quantified the most abundant flanking sequences (**Figure 2E, Supplementary Figure S7**). We found that an “AA_AA” or a “TT_TT” pattern accounted for most of the SNVs we observed, likely another reflection of the parasite’s AT-enriched genome. This pattern was most abundant around all focal nucleotides, but patterns other than these were very rare when the focal base was a C or a G. We further observed that an A or a T seemed a necessary part of both 5’- and 3’-flanking dinucleotides in all of the most abundant flanking sequences. For focal As and Ts, “AT_TA” (for focal As) and “TA_AT” (for focal Ts), i.e. patterns forming pentanucleotide sequences of alternating As and Ts, were also well represented.

### Transcriptional Sequence-Variation in *Plasmodium falciparum*

To investigate the extent of the SNVs in each *P. falciparum* line, we calculated a per-kilobase (/kb) rate of change by dividing the total number of SNVs found by the length of the *P. falciparum* genome. We obtained an average rate of ~1.6 variations/kb, i.e. ~1.6×10^−3^ variations per base. This unexpectedly high variation rate was also well-conserved between various *P. falciparum* lines, with strain- and stress-specific rates ranging from 1.47 to 1.9 variations/kb (**Figure 3**).

**Figure 3:**
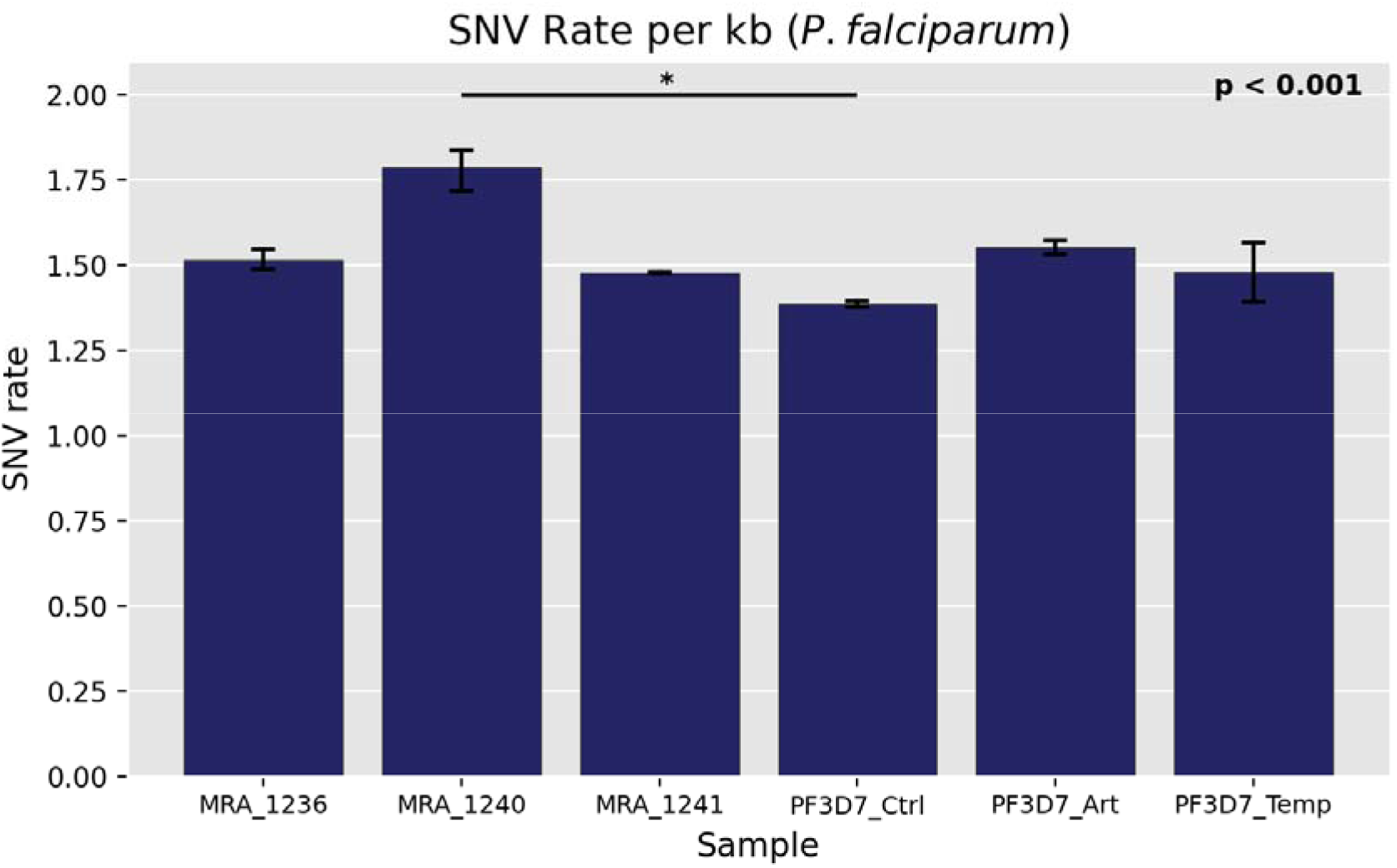
SNV rates in MRA and in-house (IH) Plasmodium falciparum lines. * denotes significance at p < 0.001 as calculated using a Dunett’s test with PF3D7_Ctrl as the control group. Error bars represent 95% confidence intervals; number of replicates = 2 (3D7_CTRL, 3D7_ART, 3D7_TEMP, MRA_1241), 3 (MRA_1236, MRA_1240).

### Transcriptional Sequence-Variation is Higher in *Plasmodium* Than Bacteria

Transcriptional sequence-level variation is often attributable to error rates associated with transcription. These rates are noted to range from ~10^−5^-10^−6^ per base in yeast^**[18]**^, similar rates in C. elegans^**[19]**^, and comparable or higher rates in bacteria ranging from 10^−4 **[2]**^ to 10^−5^-10^−6 **[1]**^ errors per base. To investigate the relative differences between the SNV rate in *Plasmodium falciparum* and other organisms, we retrieved transcriptomic data describing *E. coli, B. subtilis* [PRJNA592142], *P. vivax* (liver stages [PRJNA422240] and blood stages [PRJNA515743]), and *P. falciparum* patient isolates [PRJNA498885] from SRA and performed identical calculations to arrive at SNV rates per nucleotide and per kb using REDItools. We used filtering parameters similar to the ones described above, except for the WGS-coverage filter. We observed an average SNV rate per base of 7.23×10^−4^ for *E. coli*, 3.49×10^−4^ for *B. subtilis*, 4.35×10^−3^ for *P. vivax* schizonts+hypnozoite mixed sample, 1.80×10^−3^ for *P. vivax* hypnozoites, ~10^−3^ for *P. vivax* blood stages cultured *in vivo* in simians, and 7.42×10^−3^ in *P. falciparum* patient isolates (**Figure 4**).

**Figure 4:**
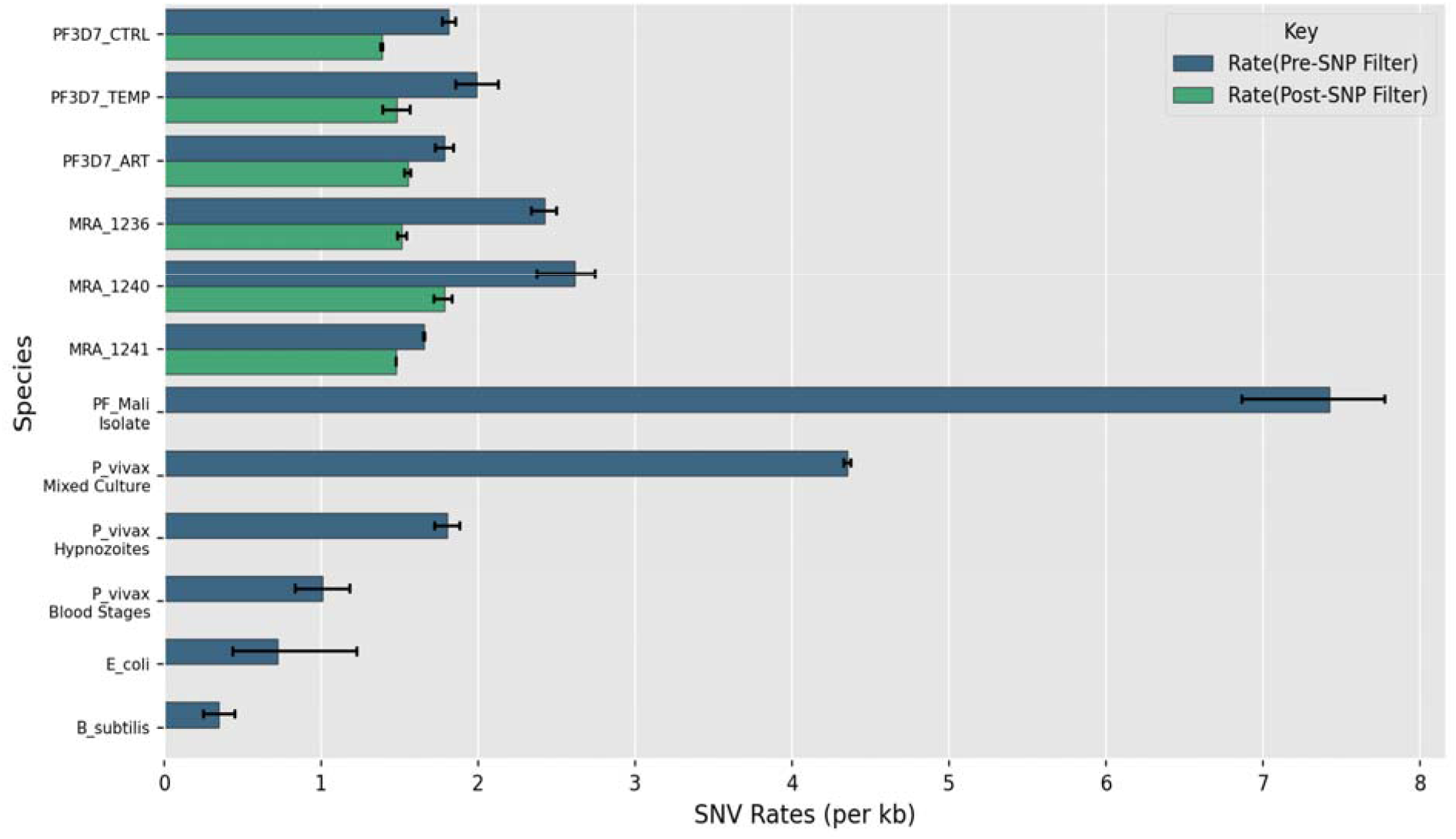
SNV rates of MR4 and in-house Plasmodium falciparum lines as compared with those for other samples and species. Error bars represent 95% confidence intervals; number of replicates = 2 (3D7_CTRL, 3D7_ART, 3D7_TEMP, MRA_1241, P. vivax Mixed Culture, P. vivax Hypnozoites), 3 (MRA_1236, MRA_1240, PF_Mali Isolate, E. coli, B. subtilis), 4 (P. vivax Blood Stages).

Since these values came from transcriptome-only analyses and SNP exclusion was not possible (this was reflected in the much higher rates we observed for the *Plasmodium spp*. samples from SRA), we also calculated the SNV rates of in-house samples without SNP exclusion (~2×10^−3^ across all sample replicates) (**Figure 4**). Interestingly, we observed that the SNV rates in *Plasmodium* species were consistently an order of magnitude higher than in bacteria, for which the reported error rates had been the highest to date (to the best of our knowledge).

## Discussion

In this work, we show that SNVs arising from base substitutions occur pervasively and non-specifically at a rate of the order of one every kilobase in the transcriptomes *P. falciparum* across treatment conditions and strains. We also show that a majority of these SNVs are predicted to have a functional impact. Given their non-specific occurrence, we speculate that these SNVs are likely reflections of RNA Pol II errors.

It is also possible that *P. falciparum* has mechanisms facilitating RNA editing, another potential source of a subset of the SNVs we report. While our search did not yield any distinct sequence motifs, nor any high-confidence RNA-editing enzymes candidates (**Supplementary Table S8**), RNA editing may still be occurring in *P. falciparum* -- the phenomenon in the parasite might be more akin to what is termed promiscuous editing^**[28]**^ in humans, wherein repetitive elements in the human transcriptome, such as Alu elements, are widely edited. Given that the *P. falciparum* genome and transcriptome are AT(/AU)-rich, it is conceivable that such a form of RNA editing may be occurring in relatively low-complexity regions of the parasite transcriptome. However, the fact that replicate datasets of any given sample showed an overlap in SNVs of no greater than ~10% indicates that RNA-editing is unlikely to be (solely) responsible for this pervasive variation.

This aforementioned AT-richness is also a characteristic of the *Plasmodium* genus, and specifically *P. falciparum* that research is still unable to fully explain. Our data shows that while A and T are the most likely to change (as would be expected from an AT-rich starting point), the probability of a focal base transitioning to any of the four bases is nearly equivalent [**Figure 2C**]. As a result, it seems that the net effect of the SNVs in *P. falciparum* is of a compensating nature, in the context of nucleotide bias. We also observe such a pattern when comparing %GC values for WGS data and the corresponding RNAseq datasets, with the %GC rising a little in the transcriptome (**Supplementary Table S10**). Previous work^**[22]**^ suggests that a large number of SNVs (as highly specific, recoding RNA editing events) in cephalopods represents a paradigm of low levels of genetic mutation, and correspondingly high levels of transcriptomic mutations, the latter providing the requisite protein diversity. In *Plasmodium*, if the numerous SNVs we report actually do lead to a nucleotide-bias compensation, then an analogous paradigm may be in play, and this may, in part, explain why the *Plasmodium* genome itself retains its AT-richness.

In the same way that the clonal *P. falciparum* cultures exhibit an inherent variation in gene expression levels^**[5,6,7,8,9]**^, our results suggest that heterogeneity at the sequence level could add a layer of complexity to the overall diversity of the parasite’s eventual phenotype. We speculate that it could be another source of variation characteristic of the parasite, conceivably arising from transcription errors, allowing a population of genetically identical cells to be phenotypically plastic to stresses or challenges. Recent work showed that the transcriptional and translational machinery of *P. falciparum* could handle the AT-richness of its genome effectively^**[29]**^. This fact, taken together with our data would seem to indicate that most of the SNVs we observed, while they might be reflections of an inherent transcriptional (i.e. RNA Pol II mediated) error rate, are likely not entirely explicable by simply the relatively lower complexity of the parasite’s transcriptome. Therefore, we anticipate that our observations, which may be generalisable for other pathogenic parasite, will lead to further investigations into the exact source(s) and consequences of the pervasive SNVs that seem characteristic of *P. falciparum*, with regard to its basic biology, possible clinical implications of such variation, and its potential interplay with the previously reported phenomena of gene-expression level variation as well as structural variations in the *Plasmodium* transcriptome.

## Materials and Methods

### Parasite Cultures

*P. falciparum* strain 3D7 was cultured as previously described^**[30]**^. Briefly, parasites were cultured in RPMI1640 medium supplemented with 25 mM HEPES, 0.5 % AlbuMAX I, 1.77 mM sodium bicarbonate, 100 μM hypoxanthine and 12.5 μg ml^−1^ gentamicin sulfate at 37 °C. Parasites were sub-cultured after every two days. Subculturing was done by splitting the flask into multiple flasks in order to maintain parasitemia around 5%. Hematocrit was maintained to 1 −1.5% by adding freshly washed O^+ve^ human RBC isolated from healthy human donors. Synchronization was done with the help of 5% sorbitol in the ring stage. Late-stage synchronization was performed using the Percoll density gradient method (63%). Parasitemia was monitored using Giemsa staining of thin blood smear.

### Stress induction

Parasites were subjected to heat and therapeutic (artemisinin treatment) stresses for 6 hours from late ring (~17 hrs) to early trophozoite (~23 hrs) stage as described earlier^**[31]**^. Briefly, double synchronization was carried out to achieve tight synchronization of parasite stages. Parasites were exposed to heat stress (40^0^C for 6 hours) and artemisinin stress (30 nM for 6 hours).

### RNA sequencing

Parasites were harvested for RNA isolation after 6 hours of stress induction. Total RNA was isolated using TRIzol reagent according to the protocol. DNAse treated RNA was used for cDNA synthesis. Quality of the RNA was verified using Agilent Bioanalyzer 2100. The cDNA libraries were prepared for samples using Illumina TruSeq RNA library preparation kit. Transcriptome sequencing was performed using Illumina NextSeq 550 system in house at IISER Pune with a standard flow cell.

### Whole genome sequencing

*Plasmodium* genome DNA was isolated using the genome DNA isolation kit. DNA concentrations were measured on the Qubit double-stranded DNA (dsDNA) HS assay kit (Invitrogen). Libraries for paired-end sequencing were constructed from DNA extracts ranging from < 50 ng/ml to 0.2 ng/µl, using the Illumina NexteraXT kit (FC-131-1024, Illumina, California, USA). The Pooled NexteraXT libraries were loaded onto an Illumina NextSeq 550 system in house at IISER, Pune with a standard flow cell.

## Data Analysis, SNV Calling and Downstream Analysis

### Quality Control and Alignment

We first checked the sequencing quality for each RNAseq and WGS sample by running each fastq file through FastQC^**[32]**^, and we trimmed the dataset to exclude positions that had a phred score of less than 25 using TrimGalore^**[33]**^ v0.6.6 (with cutadapt^**[34]**^ v3.2). We then aligned the trimmed RNAseq fastq files to the *Plasmodium falciparum* 3D7 genome (v. 41 from Ensembl) using STAR^**[35]**^ v.2.7.6a in 2-pass mode, and then indexed the resulting BAM files using samtools^**[36, 37]**^ v1.10. To align the WGS data, we used BWA (bwa mem)^**[38]**^ v0.7.17-r1198, following which we converted the resulting SAM file to a BAM file, which we sorted by coordinates and indexed using samtools. The command lines used here are provided in the Supplementary Information.

### SNV Calling and Filtering

In order to use REDItools 2.0 with python 3.6, we modified the python scripts we used to update the syntax to python 3 wherever appropriate. We then ran REDItools 2.0 using minimal command line options (**Supplementary information**) on the aligned RNAseq data and WGS data, using the RNA-table obtained from the former run as an input to the latter to specify the positions to be assayed in the WGS. The output generated by REDItools is a tab-delimited table containing data about each position read by the program in the NGS data, including the chromosome, position, reference nucleotide, coverage at that position, average read quality at that position, alternate nucleotides found at that position (if any) for the transcriptomic data (the first nine columns) and similar information for WGS data (the remaining columns). In order to annotate the RNA-table with the DNA-table, we used Annotate_with_DNA.py, which is provided as part of the REDItools suite of scripts. This script writes the WGS data for each locus in the RNA-table filling in columns 10 and further. We filtered this combined output table using *awk* command-line to exclude those positions where: (i) the WGS data showed an SNP, (ii) the frequency of base changes in RNAseq data was less than 0.1, (iii) the RNAseq data was invariant, (iv) the RNAseq coverage was less than 5 reads and (v) the WGS coverage was less than 10 reads. We then used samtools to remove duplicate reads in the aligned RNAseq data as described in Supplementary Information. Then, we reapplied REDItools to this deduplicated RNAseq file, using the previously generated, DNA-annotated and filtered RNA-table as a region file along with the previously noted REDItools options. We used the output tables from this REDItools run for further analysis.

### Downstream analysis

We used another script called AnnotateTable.py (also provided with REDItools 2.0) to annotate each SNV in the final REDItools table with gene IDs, using a sorted and trimmed gtf file (*Plasmodium falciparum* v41, from Ensembl) using the command lines (Supplementary information). Using custom python scripts, we calculated the number of SNVs occurring in each gene and the relative frequency of SNV occurrence in each gene and extracted the nucleotide sequences on either side of each changed position in order to analyse patterns in the flanking nucleotides. We also arrayed each SNV on the gene in which it occurred, dividing each gene into 100 equal-width bins, and constructed a histogram based on binning each SNV in order to analyse any positional bias of SNV occurrence on the gene body We converted the gene-annotated RNA_DNA_deduplicated file into a format resembling variant call format to use as input for snpEff, which we used according to the documentation to perform functional annotation of the SNVs in our output file. We processed the output VCF files generated by snpEff using snpSift to filter the file and retain data related to predicted amino-acid changes. We analysed this filtered VCF file using a custom script to visualise the spectrum of amino-acid changes in each sample. We did not retain any annotations of the types “upstream variant” and “downstream variant”, since snpEff defines upstream and downstream regions as 5 kilobases in length, which is too large for the relatively compact genome of *P. falciparum*. SNV rates per base were calculated as the number of SNVs in the filtered output divided by the genome length in nucleotides. We used Salmon^**[39]**^ to obtain gene abundance estimates in transcripts per million.

### Analysis of Samples Sourced from SRA

NGS (RNAseq) datasets for *Plasmodium falciparum* patient-isolates from Mali, *Plasmodium vivax* liver-stages (mixed cultures and hypnozoites), *Plasmodium vivax* blood-stages, *Escherichia coli*, and *Bacillus subtilis* were downloaded using SRA toolkit^**[40]**^ as fastq files. We used genome versions **PvP01** for *P. vivax*, **ASM584v2** for E. coli, and **ASM608879v1** for *B. subtilis*. The bacteria datasets were aligned using BWA, while the *Plasmodium* spp. datasets were aligned using STAR. We analysed each dataset with REDItools 2.0, filtering the output files to remove positions where; (i) the frequency of base changes in RNAseq data was less than 0.1, (ii) the RNAseq data was invariant, (iii) the RNAseq coverage was less than 5 reads and (iv) The average phred-score was less than 25. SNV rates per base were calculated as the number of SNVs in the filtered output divided by the genome length in nucleotides.

### Circos plot generation

We used the R package RCircos^**[41]**^ to generate circos representations for the SNV distribution on the whole-genome scale.

### Statistics and Error Bars

A Pearson’s Chi-squared test was performed to assess whether the distribution of frequency of SNV types (by base-substitution, as % of total) was significantly different from a uniform distribution (with the assumption that all base-substitutions are equally likely to occur). The test was performed in R using the chisq.test() function A Dunnett’s test was used to calculate statistical significance for the differences in average SNV rates between the 3D7 parasite cultures and the three MRA parasite lines. This test was performed in R using the DunnettTest() function from the R library DescTools, using PF3D7_Ctrl as the control set. For Figures 2, 3 and 4, error bars indicate 95% confidence interval for replicates and the number of replicates for each sample is as follows:

- *P. falciparum* MRA_1236: 3 replicates
- *P. falciparum* MRA_1240: 3 replicates
- *P. falciparum* MRA_1241: 2 replicates
- *P. falciparum* 3D7 Control: 2 replicates
- *P. falciparum* 3D7 Drug-stressed: 2 replicates
- *P. falciparum* 3D7 Temperature-stressed: 2 replicates
- *P. falciparum* Mali Isolates [PRJNA498885]: 3 replicates
- *P. vivax* [PRJNA422240] (mixed culture): 2 replicates
- *P. vivax* [PRJNA422240] (hypnozoites only): 2 replicates
- *P. vivax* [PRJNA515743]: 4 replicates
- *E. coli* [PRJNA592142]: 3 replicates
- *B. subtilis* [PRJNA592142]: 3 replicates

## Code and Data Availability

All custom code written for the analysis described here will be made available on request via GitHub.

## Acknowledgements

We thank Prof. Sutirth Dey for his valuable inputs and suggestions on this work. The following reagents were obtained through BEI Resources (www.mr4.org), NIAID, NIH: *Plasmodium falciparum*, Strain IPC 3445 (MRA-1236), Strain IPC 5202 (MRA-1240), Strain IPC 4912 (MRA-1241), contributed by Didier Menard. This work was supported by DBT-Genome Engineering Technologies program (BT/PR25858/GET/119/169/2017) from the Government of India to KK. The funders had no role in study design, data collection and analysis, decision to publish, or preparation of the manuscript.

## Authors’ Contributions

BD designed, performed experiments, and analyzed data. AK and MDV cultured *P. falciparum* and generated NGS data. BD and KK wrote the manuscript. KK planned, coordinated, and supervised the project. All authors read and approved the final manuscript

