## Supplemental Figures for "Exceptionally high sequence-level variation in the transcriptome of *Plasmodium falciparum*"

**Exceptionally high transcriptional error rates in *Plasmodium falciparum***

**Supplementary Figures and Tables**

All supplementary tables are provided in a separate file.

**Supplementary Table S1** contains a comprehensive overview of the transcriptome data presented in this work, including sample names, sources of data, the number of SNV positions and the SNV rate (per kilobase) calculated for each sample before and after filtering & downstream analysis. In both cases, the columns containing the descriptor “Post-Processing” indicate that the values noted there have been calculated after the annotation of the REDItools RNAtable with the corresponding DNAtable, the filtering of this merged table, and the second round of REDItools performed using deduplicated BAM files and this merged table. STable1 also contains the sequencing and alignment properties of each sample and replicate -- it contains the average read length, the number of input reads, the % of reads aligned and unaligned, the number of reads aligned, the genome length (in base pairs) used for SNV rate calculation and sequencing depth calculation, and “genome coverage”, an ad hoc value representing sequencing depth of aligned reads, calculated as the number of nucleotides aligned in each sample divided by the genome length. Alignment statistics for all samples except bacteria are from the respective STAR Log.final.out files; statistics for the bacterial transcriptome were extracted from BWA-aligned output using samtools.

**Supplementary Table S2** records the number of genes which were affected by SNVs for each sample. In order to arrive at the frequency, RNA coverage and DNA coverage cutoff values for the merged REDItools table, we used a range of cutoffs and plotted the number of values retained after filtering with that cutoff. For DNA coverage cutoff estimation, the Frequency and RNA coverage cutoffs were kept at 0.1 and 5 respectively **(Supplementary Figure S1, Supplementary Table S3)**. For the RNA coverage cutoff estimation, Frequency and DNA coverage cutoffs were kept at 0.1 and 10 respectively **(Supplementary Figure S2, Supplementary Table S4)**. For frequency cutoff estimation, the DNA coverage and RNA coverage cutoffs were kept at 10 and 5 respectively **(Supplementary Figure S3, Supplementary Table S5).**

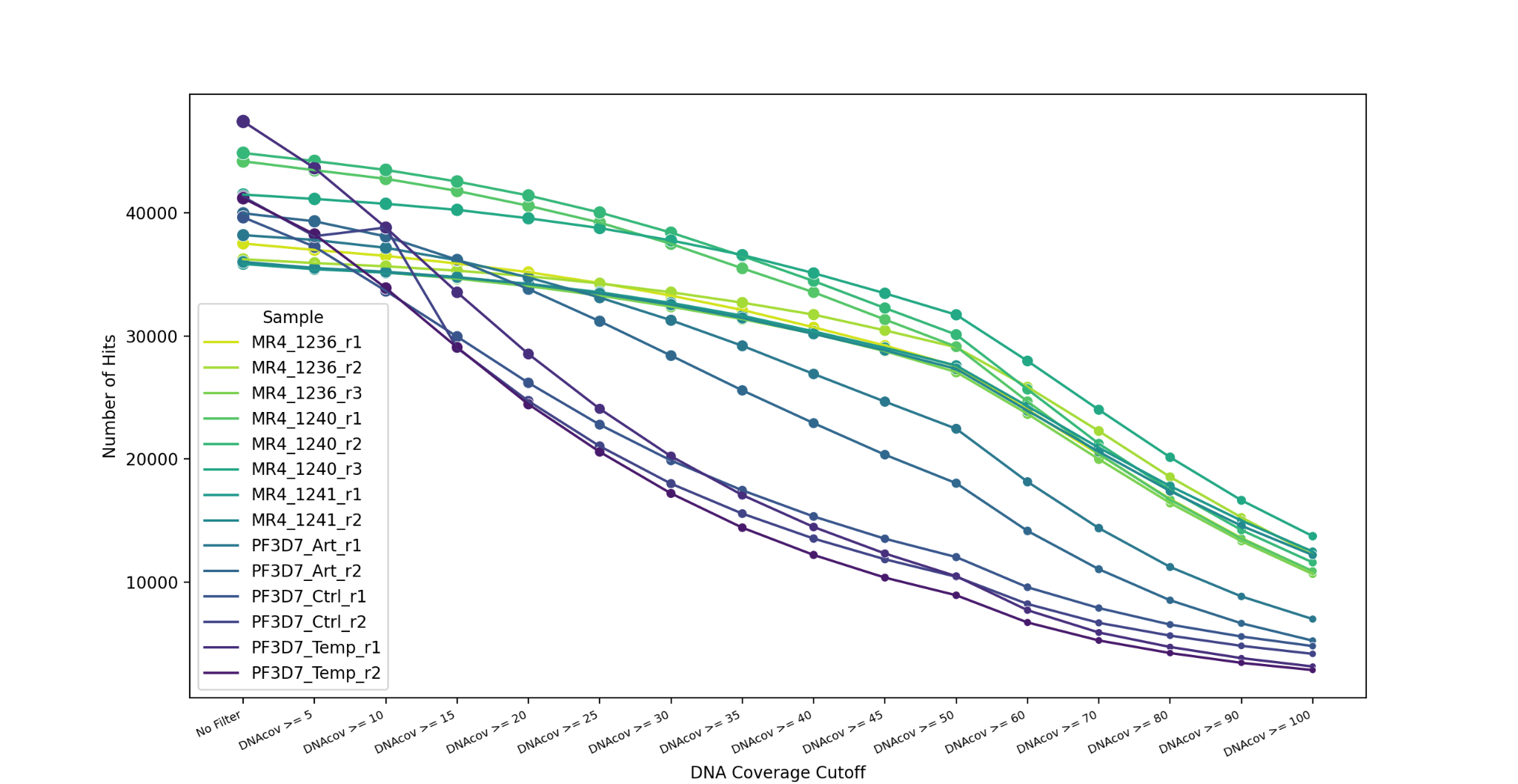

*Supplementary Figure S1: Number of Hits Retained at Varying DNA coverage Cutoffs. “No Filter” on the X-axis represents a dataset from which genomic variations have been removed but no further filter has been applied*

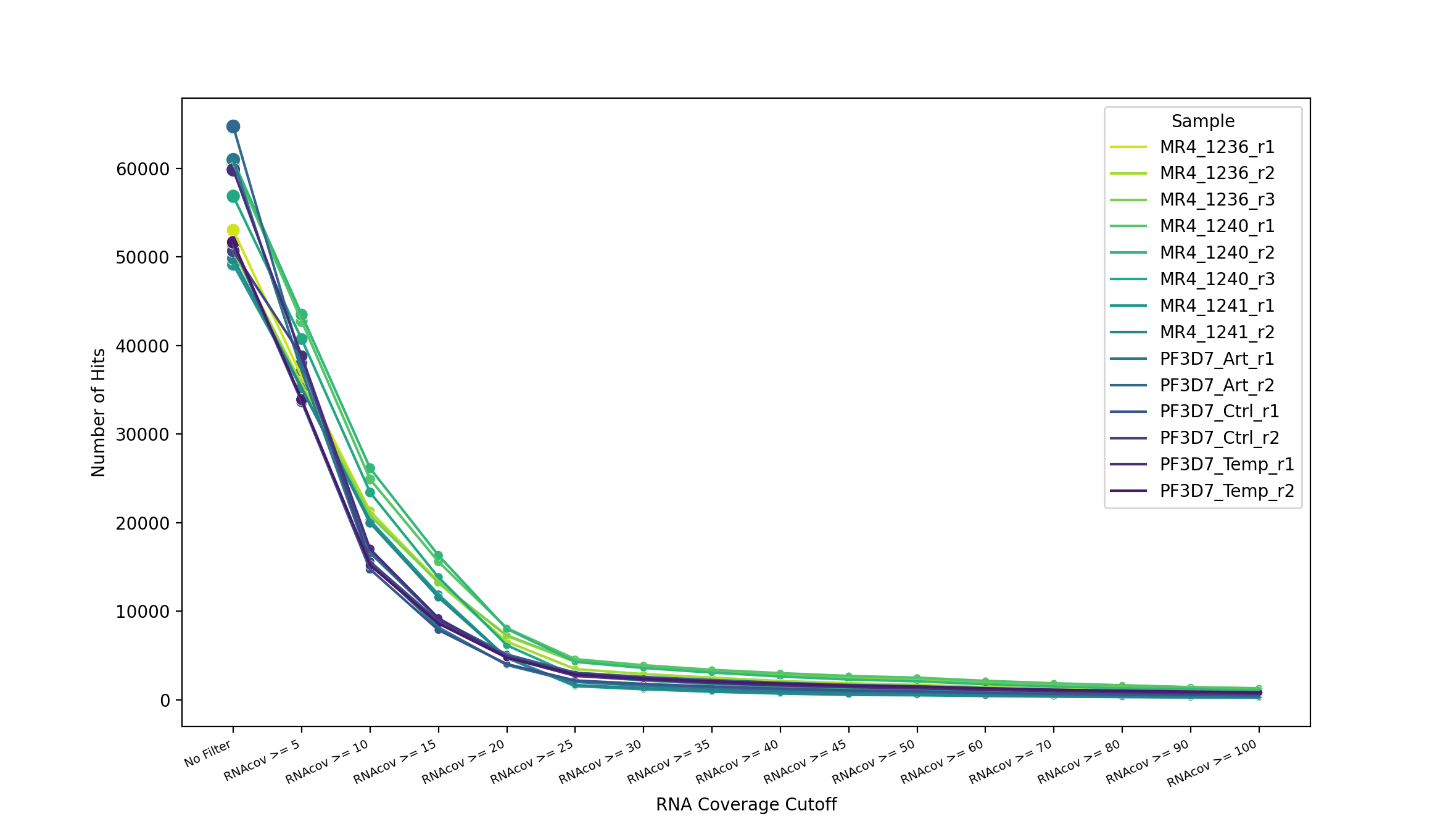

*Supplementary Figure S2: Number of Hits Retained at Varying RNA coverage Cutoffs. “No Filter” on the X-axis represents a dataset from which genomic variations have been removed but no further filter has been applied*

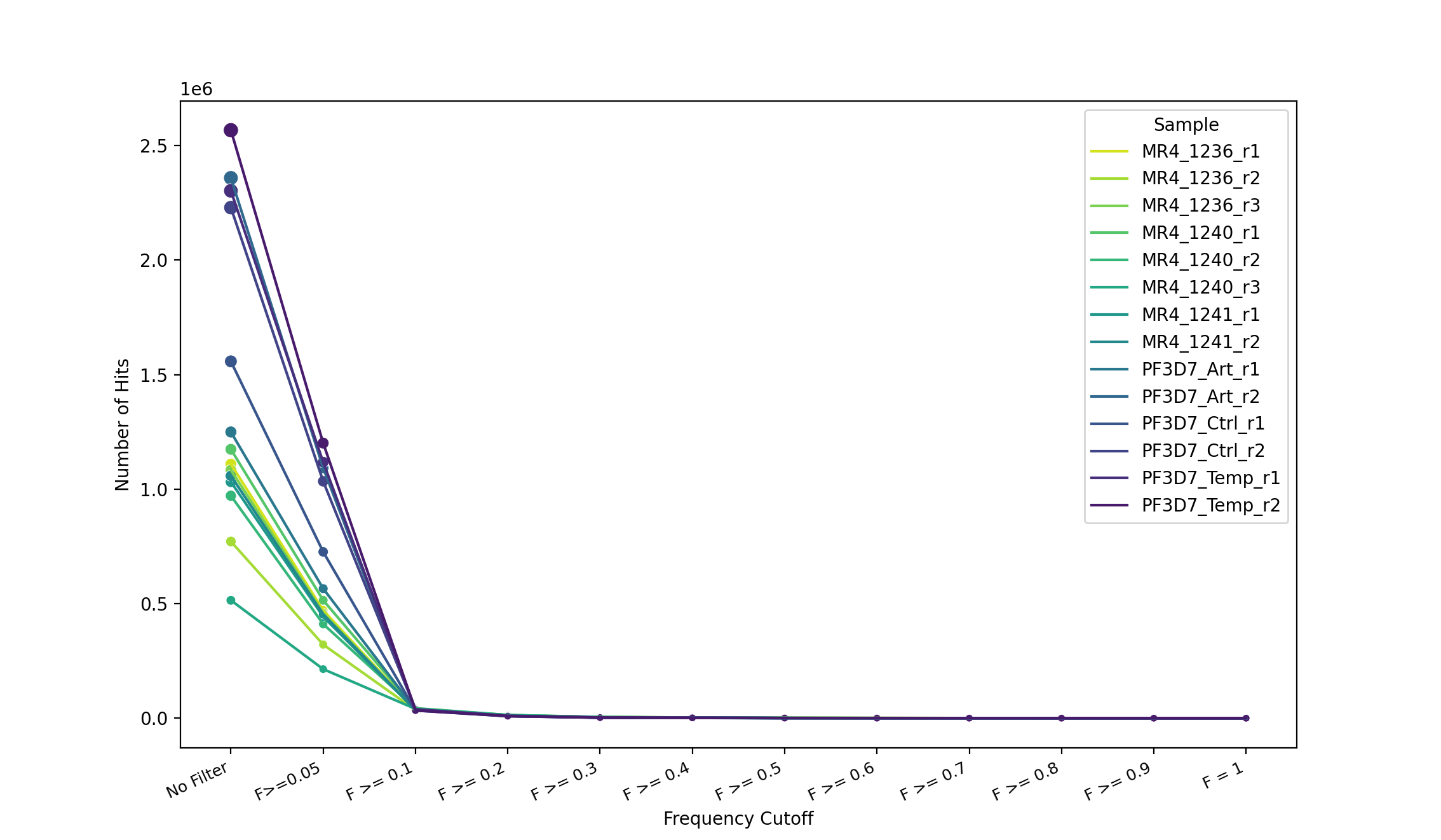

*Supplementary Figure S3: Number of Hits Retained at Varying Frequency Cutoffs. “No Filter” on the X-axis represents a dataset from which genomic variations have been removed but no further filter has been applied*

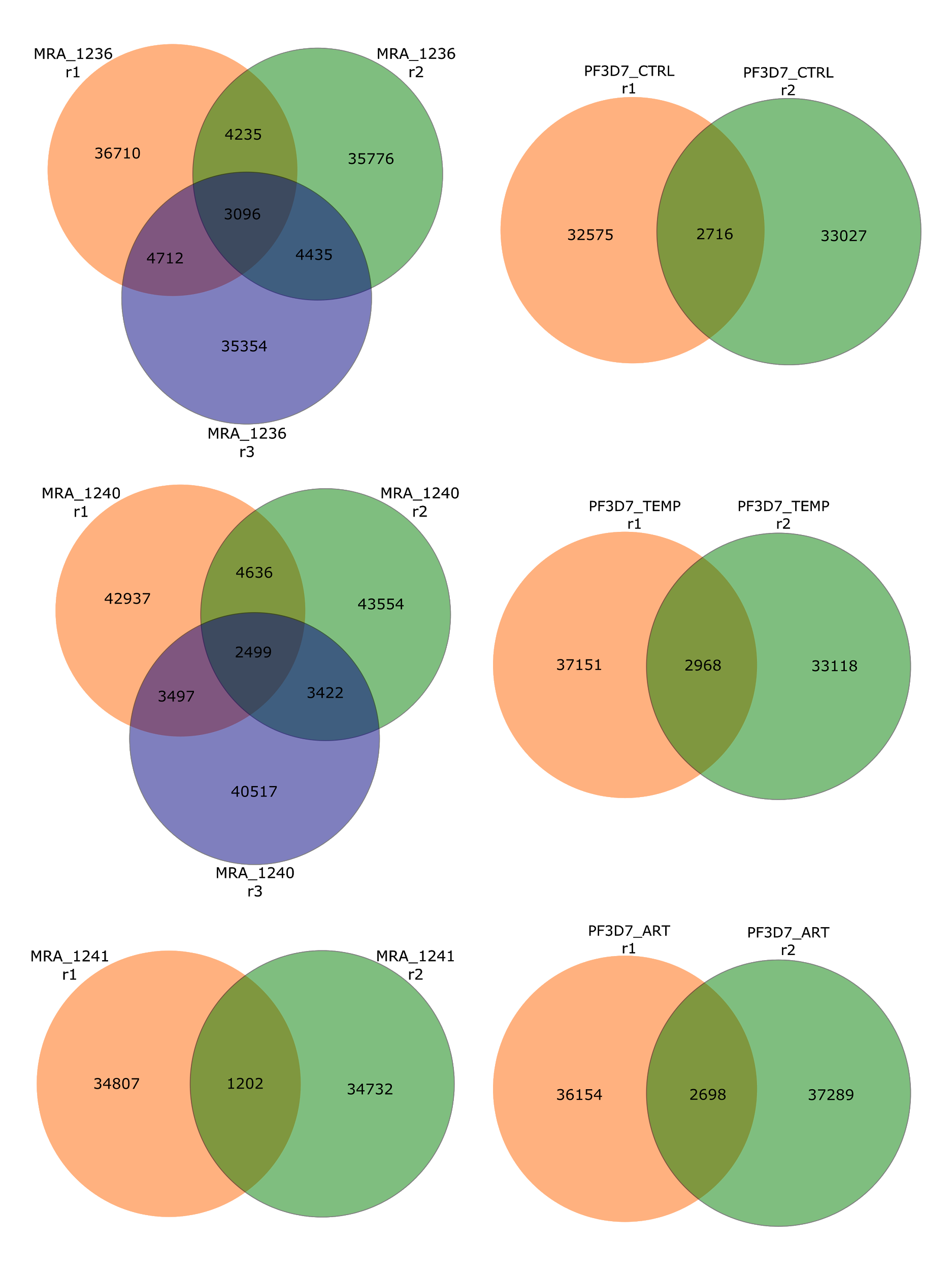

*Supplementary Figure S4: Extent of Overlap Between Replicates. Samples labels are outside the Venn diagram; each circle represents one replicate; the total number of SNVs post-filtering are noted for each replicate and numbers in overlaps represent the number of SNV positions common between replicates. An SNV is considered to be common between replicates if it is located in the same position AND shows the same type of base substitution at that position (i.e. if the SNV location AND type are the same between replicates)*

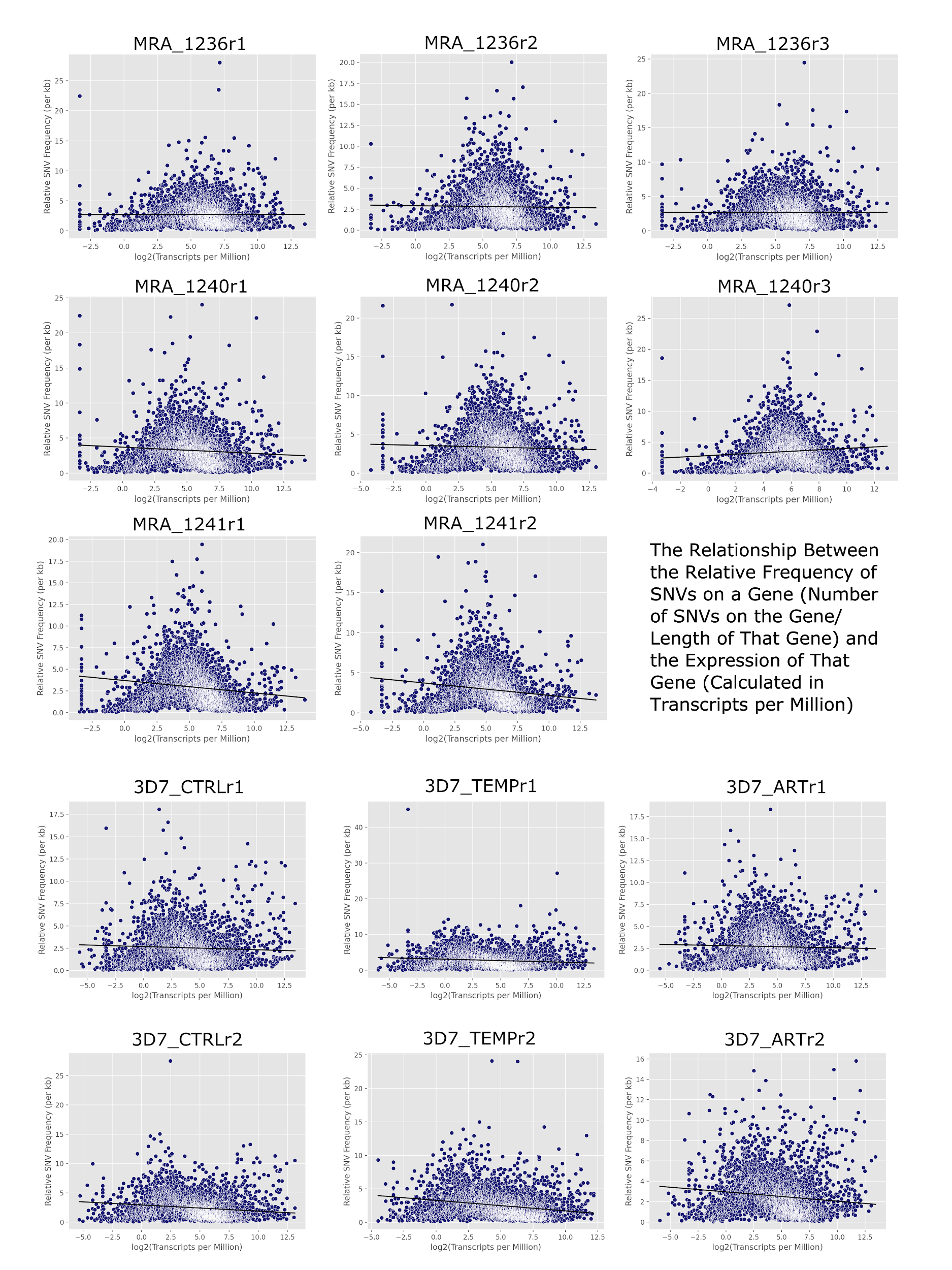

*Supplementary Figure S5: Relationship Between the Relative Frequency of SNVs in a Gene (per kb) and Its Expression Level in log2(Transcripts per Million). Black lines are correlation trend lines. With the exception of MRA_1240r3, relative SNV frequencies (defined as the number of SNVs in a gene divided by the length of that gene) for all samples show a weakly negative or no correlation to the expression levels of the corresponding genes.*

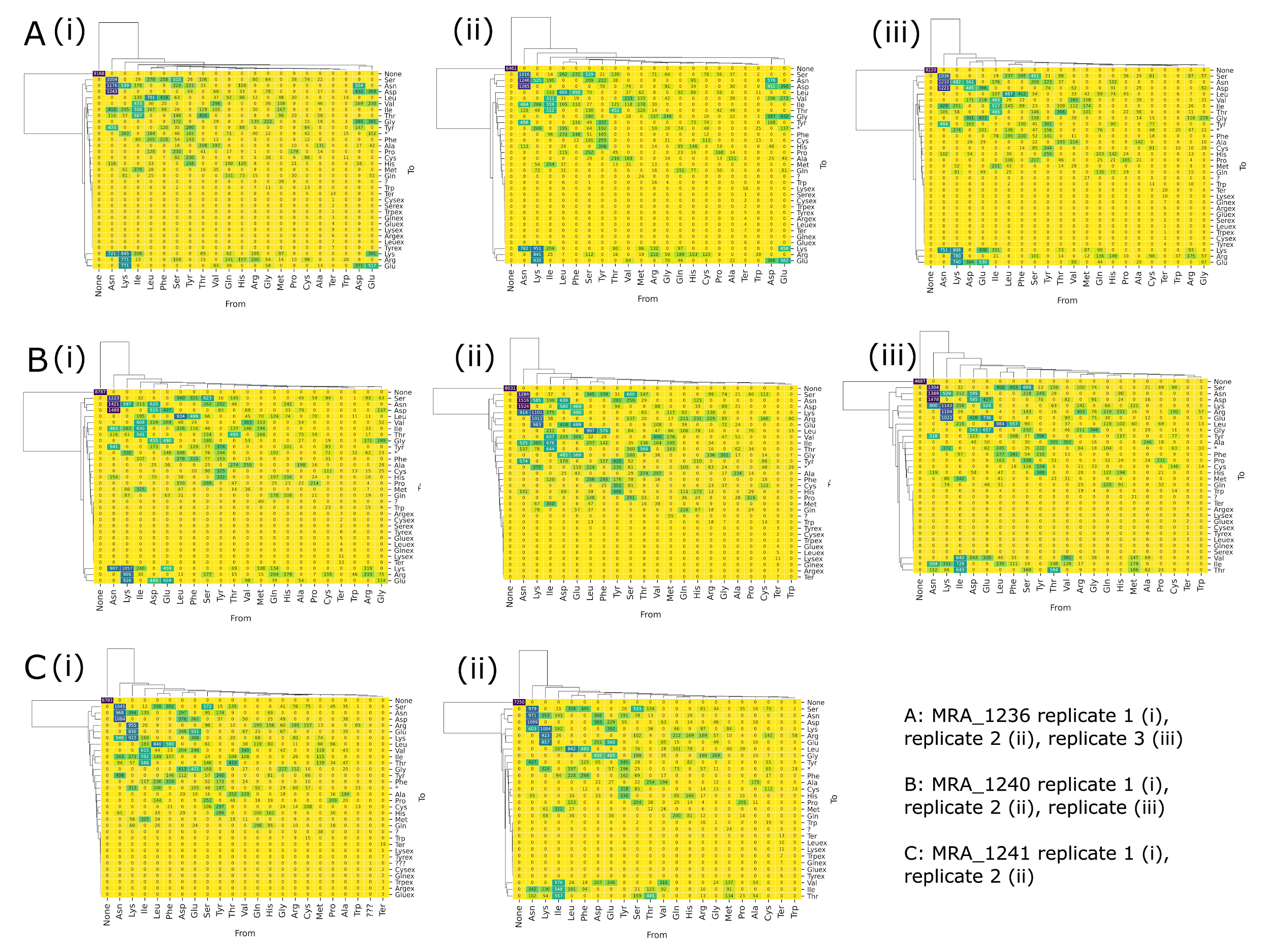

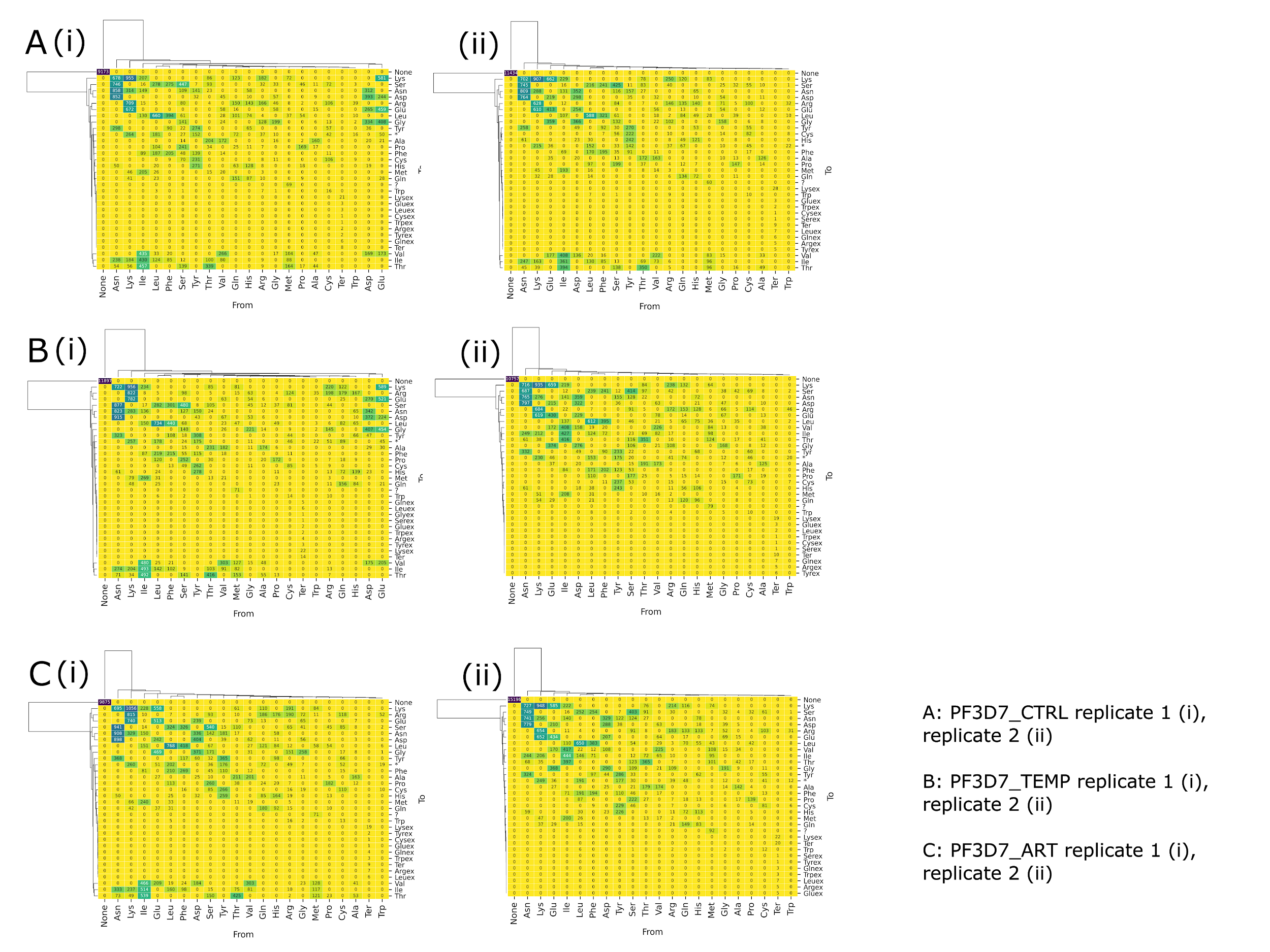

*Supplementary Figure S6: Spectrum of Amino Acid Changes in MRA and PF3D7 Lines. Darker colours indicate higher frequency; heatmaps record the frequency of amino acid changes from amino acids on the horizontal axis to those on the vertical axis.*

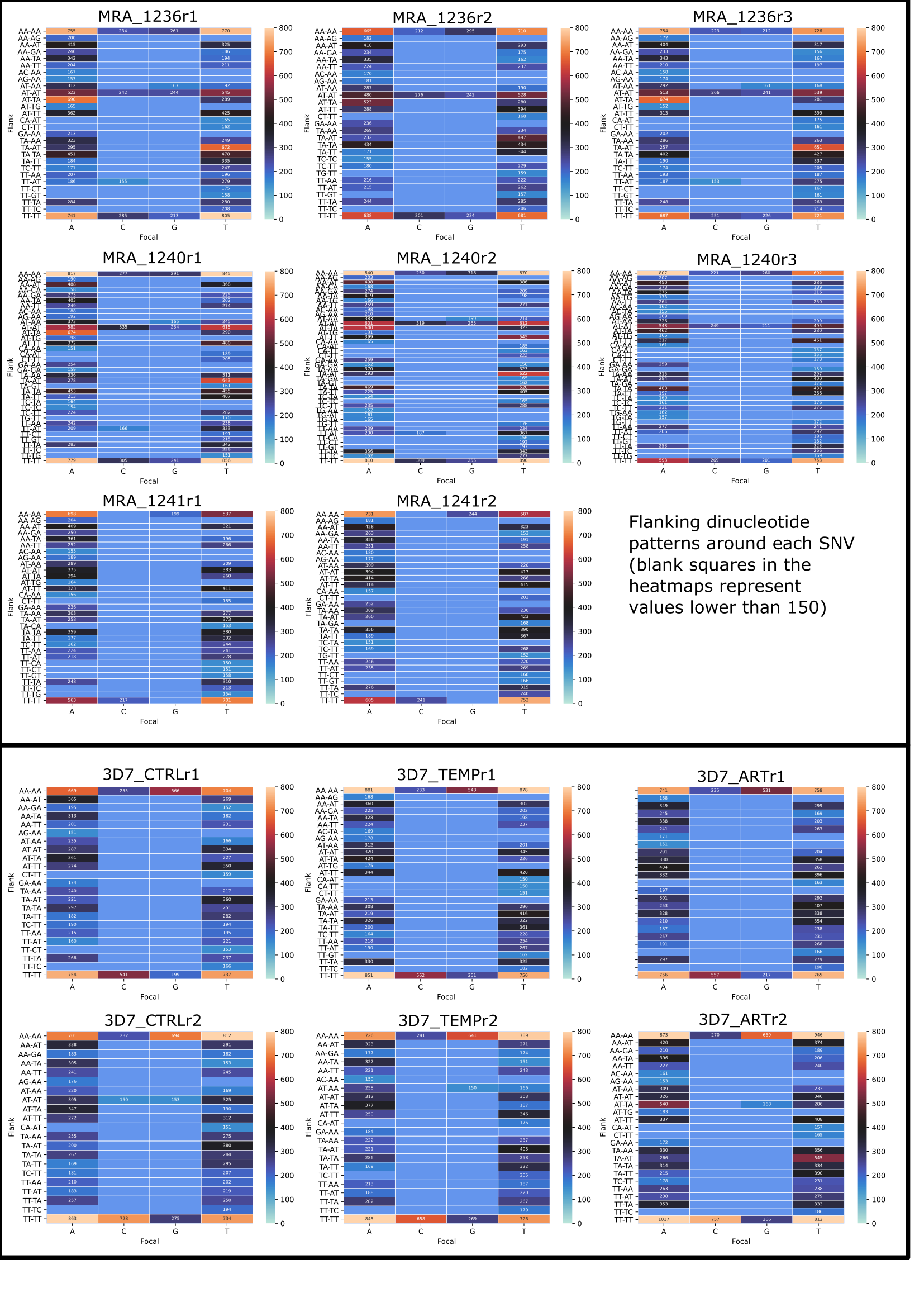

*Supplementary Figure S7: Abundant Dinucleotide Patterns Flanking SNV Positions. The vertical axis shows the most abundant flanking dinucleotides around each SNV position, where the reference/focal base is recorded on the horizontal axis. Empty panels in the heatmaps indicate frequencies of occurrence of less than 150.*

**Supplementary Table S6** contains data about hits corresponding to each type of base change, where X > Y indicates nucleotide X changing to nucleotide Y. **Table S6.1** shows the number of hits for each X > Y combination and **Table S6.2** shows the proportion of each change type as percentage of each type out of the total.

**Supplementary Table S7** describes base change patterns, where X> nucleotide X changing to another nucleotide, and >X indicates that a reference nucleotide changed to nucleotide X. **Table S7.1** records the number of hits corresponding to each shift type and **Table S7.2** records the same data as percentages of total.

**Supplementary Table S8** describes the results of a BLAST based search for potential RNA-editing enzymes, using known RNA-editing enzymes from *H. sapiens* and *T. brucei* as query sequences. Column 1: the source organism of the BLAST query sequence; column 2: gene name associated with query sequence; column 3: domain of protein used as query sequence (sequence start index-sequence end index); column 4: gene names of *P. falciparum* genes matching the query sequence; column 5: annotated function of the matches recorded in *P. falciparum* (source of annotation data: PlasmoDB); column 6: BLAST match score; column 7: E-value of the match; column 8: gene ontology terms relevant to putative RNA editing activity (if any).

**Supplementary Table S9** records the number of SNVs annotated to a specific predicted functional effect. “Intronic” indicates that the SNV was predicted to be in an intronic region, “Non-coding” indicates that the SNV occurred in a non-coding region of a transcript, “Missense” indicates that the SNV caused a non-synonymous amino acid change, “Start_lost” indicates that the SNV resulted in a start codon being changed, “Stop_gained” indicates that the SNV led to a premature stop codon forming, “Stop_lost” indicates the mutation of a stop codon to an amino acid codon, “Splicing” indicates that the SNV caused a splicing variant, “Synonymous” indicates a mutation of the reference amino acid into a synonymous one, “ncRNA Exon” indicates that the SNV occurred in a non-coding transcript. **Table S9.1** records the number of SNVs associated with each functional prediction and **Table S9.2** records the same data as the percentage total.

**Supplementary Table S10** shows the GC content of the quality-trimmed fastq files for each WGS sample and the corresponding RNAseq replicates. In the table, %GC for a sample is calculated as the average %GC of the two fastq files (forward and reverse) for each dataset; the values used for this calculation are as reported in the FastQC summary html file.
